# Whole-brain causal connectivity during decoded neurofeedback: a meta-study

**DOI:** 10.1101/2024.11.16.623939

**Authors:** Fahimeh Arab, AmirEmad Ghassami, Hamidreza Jamalabadi, Megan A. K. Peters, Erfan Nozari

## Abstract

Decoded Neurofeedback (DecNef) enables modulation of targeted brain activity patterns without subjective awareness through multivariate pattern analysis, reinforcement learning, and real-time fMRI feedback. Despite its proven effectiveness, the causal mechanisms underlying DecNef and the neural dynamics that distinguish successful learners from those who struggle remain poorly understood. We conducted a meta-study using fMRI data from five DecNef experiments and extracted causal network dynamics as well as their associations with performance differences. Across studies, we found that connectivity within a posterior control hub–consisting of posterior cingulate, precuneus, and lateral posterior parietal cortices–is stronger during DecNef and positively correlates with neurofeedback success. Comparisons across cognition- and perception-targeted DecNef revealed separation in connections to somatomotor network, where connections between somatomotor and control-default-attention networks are larger during cognitive neurofeedback while connections between somatomotor and subcortical-visual-limbic networks are larger during perceptive DecNef. Whole-brain causal connectivity during DecNef further exhibited distinct network reorganizations, with greater subject-to-subject variability, increased engagement of control, limbic and visual, and decreased engagement of ventral and dorsal attention networks. Our results distill the complex and distributed network mechanisms underlying DecNef into dissociable roles for well-known functional subnetworks, thus advancing the research and clinical applications of decoded neurofeedback.

## Introduction

Two decades have passed since Weiskopf et al. (2004)’s pioneering demonstration of the feasibility of using real-time fMRI as a brain-computer interface, enabling participants to self-regulate brain activity via feedback. More recently, decoded neurofeedback (DecNef) has been proposed as a novel technique combining implicit neurofeedback and multivariate pattern analysis (Shibata et al., 2011; Taschereau-Dumouchel et al., 2021). Unlike traditional methods that rely on overall signal amplitude and explicit strategies, DecNef induces specific signal patterns in target brain regions, altering these neural patterns (and subsequently impacting behavior) without participants’ awareness of the exact content and purpose of the manipulation (Cortese et al., 2021; Shibata et al., 2019, 2011). As a result, DecNef can help reduce potential confounding effects from cognitive processes or awareness of the specific dimension being manipulated (Cortese et al., 2021). These characteristics have made DecNef especially well-suited for developing new clinical applications, particularly in the treatment of neuropsychiatric disorders (Chiba et al., 2019; Koizumi et al., 2016; Taschereau-Dumouchel et al., 2018, 2020; Yamada et al., 2017). DecNef has also proved valuable beyond clinical applications, offering insights in systems and cognitive neuroscience to explore fundamental brain functions in diverse areas such as visual sensitivity (Shibata et al., 2011), color perception (Amano et al., 2016), fear memory (Koizumi et al., 2016; Taschereau-Dumouchel et al., 2018), facial preference (Shibata et al., 2016), and perceptual confidence (Cortese et al., 2016).

The precise neural mechanisms underlying DecNef are nevertheless poorly understood. In particular, a major question in neurofeedback research is understanding the factors that contribute to successful self-regulation. Recent research has started to delve into these questions through a variety of methods, including meta-analyses, computational models, and neural network simulations (asr, 2024; Emmert et al., 2016; Haugg et al., 2020; Oblak et al., 2017, 2019; Pereira et al., 2024; Sepulveda et al., 2016; Shibata et al., 2019; Skottnik et al., 2019). One plausible mechanism that has been suggested is reinforcement learning (Lubianiker et al., 2022). In particular, Shibata et al. (2019)’s *targeted neural plasticity model* suggests that DecNef likely drives neural plasticity through reinforcement learning mechanisms, with significant activation in reward-related brain regions such as the ventral striatum and putamen in response to feedback signals. This suggests that DecNef engages the brain’s reward-processing circuits and may share neural foundations with conventional neurofeedback and brain-machine interfaces. Yet, existing models do not address the network dynamics that give rise to the induction process on a trial by trial basis and what network processes distinguish successful learners from those who struggle. Our work addresses this gap by identifying whole-brain causal interactions during DecNef induction sessions compared to baseline. By examining causal connectivity patterns, we provide a connectivity-based understanding of DecNef induction dynamics and offer insights into the hierarchical mechanisms that govern decoded neurofeedback.

Our work draws from the extensive literature of causal discovery (Assaad et al., 2022; Glymour et al., 2019) to extract causal brain mechanisms from purely observational fMRI. fMRI possesses a major advantage for causal discovery because of its potential for whole-brain coverage, but it also poses significant challenges. fMRI’s low temporal resolution, combined with the computational complexity of analyzing large-scale networks, makes it difficult to accurately discern directional relationships between brain regions. Traditional methods such as Granger Causality (GC)(Barnett and Seth, 2014; Granger, 1969) and Dynamic Causal Modeling (DCM) (Friston et al., 2014) often struggle to handle these complexities (Arab et al., 2025). As such, in previous work we have developed the Causal discovery for Large-scale Low-resolution Time-series with Feedback (CaLLTiF) algorithm to address these challenges and learn causal graphs based on both lagged (slow) and contemporaneous (fast) relationships (Arab et al., 2025). When applied to synthetic fMRI data, CaLLTiF outperformed a wide range of state-of-the-art methods in terms of accuracy and scalability. When applied to resting-state human fMRI, CaLLTiF uncovered causal connectomes that were highly consistent across individuals and showed a top-down causal flow from attention and default mode networks to sensorimotor regions, Euclidean distance-dependence in causal interactions, and a strong dominance of contemporaneous effects.

Building on these insights, in the present study we used CaLLTiF to learn causal graphs from fMRI data of DecNef induction sessions and compared them to causal graphs learned from baseline fMRI of respective subjects. We conducted a meta-study across five previously-published DecNef experiments, each involving a different induction target pattern and multiple fMRI sessions per participant. This meta analysis makes it possible to move beyond the specifics of each task and identify core causal mechanisms that are shared across all tasks and subjects (n = 45).

## Results

### Causal discovery from decoded neurofeedback

Figure 1a illustrates the general framework of Decoded Neurofeedback (DecNef), consisting of decoder construction (DC) and neurofeedback (NF) sessions (Cortese et al., 2021). In DC sessions, multivariate pattern analysis (MVPA) is used to train a decoder (e.g., logistic regression) on BOLD activations in a select number of voxels. In NF sessions, this decoder is applied in real time and its output is shown in the form of visual feedback to the participants, which they then use to self-regulate the specific voxel activation pattern sought by the decoder. In this work, we use combined data from five DecNef studies, made publicly available by Cortese et al. (2021).

**Figure 1.**
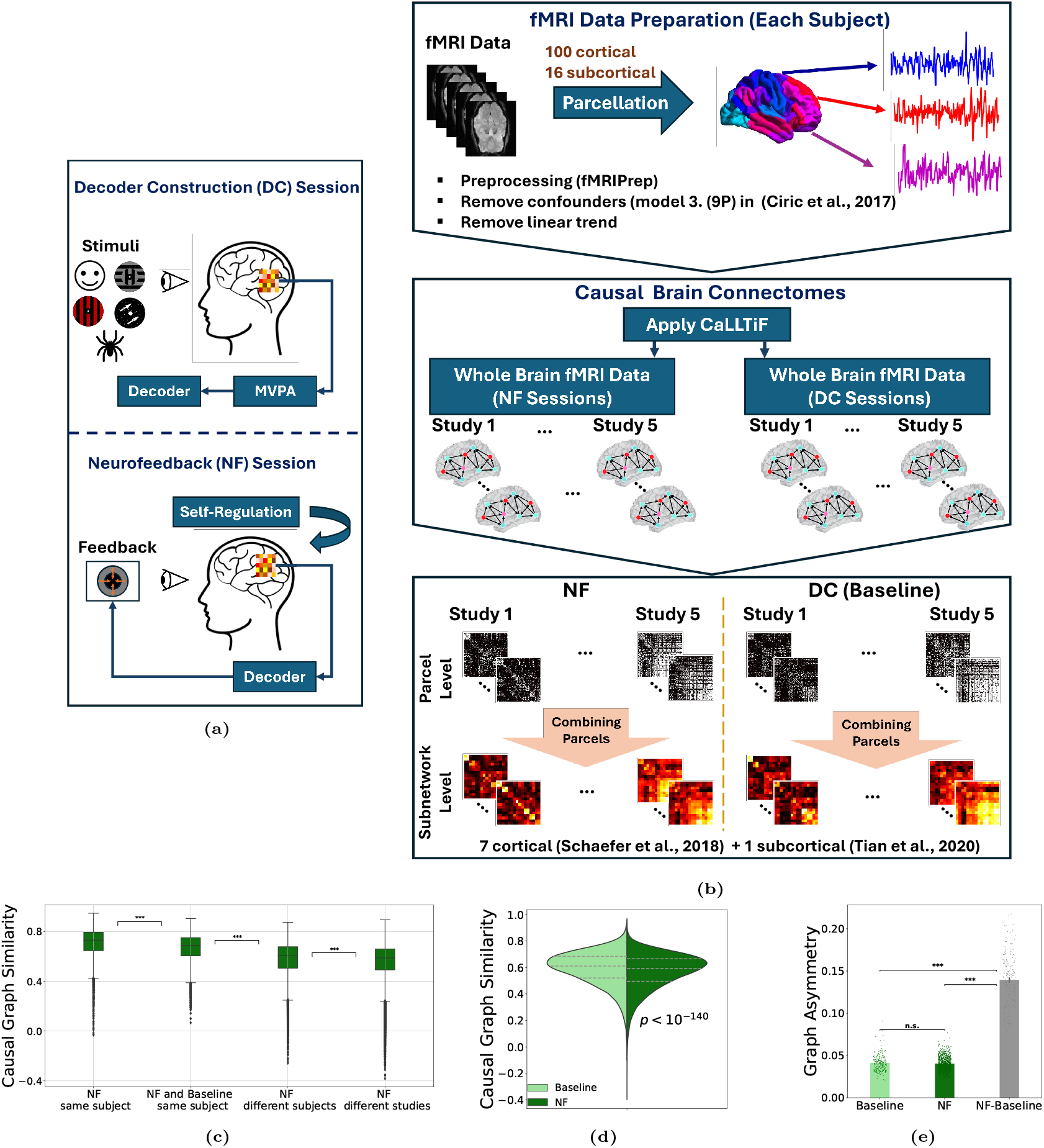
Overview of the decoded neurofeedback (DecNef), data processing, and causal discovery pipelines. **(a)** During the Decoder Construction (DC) sessions (baseline), participants viewed stimuli while their whole-brain fMRI was recorded and used offline to construct decoders through multivariate pattern analysis (MVPA). In the Neurofeedback (NF) sessions, participants engaged in self-regulation of brain activity, guided by real-time feedback based on decoders trained on data from the DC sessions. **(b)** For each subject, fMRI data was preprocessed using fMRIPrep (skull stripping, motion correction, spatial normalization, and smoothing) followed by 9P confound removal and linear detrending. Data were then parcellated into 100 cortical regions (Schaefer 100x7 atlas (Schaefer et al., 2018)) and 16 subcortical ones (Melbourne Scale I atlas (Tian et al., 2020)). We then applied CaLLTiF to fMRI data from both NF and DC sessions of each subject to generate causal connectivity graphs. Parcel-level graphs were finally summarized into subnetwork-level graphs, each consisting of 16 nodes across both hemispheres (see Methods). **(c)** Hierarchy of similarity of causal graphs across conditions. Similarity was measured via Pearson correlation coefficient between pairs of vectorized subntwork-level graphs. **(d)** Similar to (c) but for similarity across all NF and baseline graphs. NF graphs exhibit greater variability across subjects and sessions compared to baseline. Each distribution shows pairwise graph similarities combined across all studies, subjects, sessions, and runs. **(e)** Distribution of asymmetry of causal graphs during baseline, NF, and NF−baseline. For each graph with weighted connectivity matrix *A*, asymmetry is computed as ∥*A* − *A*^*T*^ ∥_1_*/*∥*A* + *A*^*T*^ ∥_1_.

Figure 1b shows the pipeline for the causality analysis conducted in this paper. Using our recently proposed algorithm CaLLTiF (Arab et al., 2025), we derived causal brain connectomes from whole-brain fMRI data collected during both NF and DC sessions, with the latter serving as a subject- and task-specific baseline for NF sessions. Before running CaLLTiF, fMRI data was preprocessed and parcellated into 100 cortical (Schaefer et al., 2018) and 16 subcortical (Yeo et al., 2011) regions (see Methods). We then applied CaLLTiF separately to data from each session, resulting in (study, subject, session)-specific parcel-level directed causal graphs (see Methods). Parcel-level graphs were then aggregated to obtain subnetwork-level graphs in which each node is a “functional network” (Yeo et al., 2011). The latter graphs consist of 16 nodes—7 cortical (Schaefer et al., 2018) and 1 subcortical (Tian et al., 2020) separately across the left and right hemispheres. We then statistically compared subnetwork-level graphs between NF and baseline sessions to identify core causal mechanisms underlying DecNef that are shared across all subjects and studies.

### Causal connectivity during NF is highly stable and personalized

We first examined the similarity of causal graphs across sessions, subjects, and studies, measured by Pearson correlation coefficients between pairs of subnetwork-level graphs (FDR-corrected *p <* 0.05, Student’s *t*-test). As seen from Figure 1c, graph similarities are generally high and follow an expected hierarchy, with the strongest similarities existing between NF sessions of the same subjects and least similarity (greatest variability) seen across causal graphs of different studies. This suggests that CaLLTiF-derived causal graphs are generally stable and individualized, with NF sessions introducing repeatable causal patterns within each subject that set them apart from baseline sessions. Across all subjects and studies, however, NF graphs are *less* similar than baseline graphs (Figure 1d). This distinction highlights the adaptive and personalized nature of NF, where each subject’s unique self-regulation strategies contribute to greater variability compared to baseline sessions that follow a relatively consistent task design across all studies.

Further, our results show that causal graphs are more similar between NF and baseline sessions of the same subject than they are between NF sessions of different subjects, even within the same study (Figure 1c). This highlights the importance of subtracting baseline graphs for each individual in order to control for strong subject-to-subject variability that exists independently of the NF process. Notably, difference graphs (NF - baseline) are also significantly more directed than both baseline and NF graphs, underscoring the subtraction of symmetric baseline connectivity in order to better distinguish the directions of causal flow underlying NF regulation (Figure 1e). Finally, note the lowest similarity observed between NF graphs across different studies (Figure 1c), confirming the presence of unique causal patterns driven by study-specific protocols or task demands. In most of the ensuing analyses we seek to “average-out” these study-specific effects and extract shared causal effects that underlie DecNef, independently of the NF’s target area or pattern.

### Causal connectivity during NF involves increased engagement of control, limbic, and visual networks and decreased involvement of attention networks

We next examined the strengths of identified causal edges, at the subnetwork level, during NF and baseline. Both NF and baseline graphs predominantly exhibited excitatory connections (Supplementary Figure 2). Compared to baseline, each causal edge during NF, whether excitatory (positive) or inhibitory (negative), belongs to one of two general categories: those that increased in magnitude during NF (Figure 2a,b,e), and those that became weaker during NF (Figure 2c,d,f). While most nodes have edges in both categories, a few nodes stand out as having disproportionately more edges in one category or another. In particular, the vast majority of casual edges connected to control, limbic, and visual networks became stronger during NF compared to baseline, indicating an overall increased engagement of these networks during NF. In contrast, most edges connecting attention networks to the rest of the brain became weaker, marking a reduced involvement of attention networks during NF. The remaining networks (default mode, somatomotor, and subcortical) involve a comparable mix of edges that became stronger and those that became weaker, leaving their overall involvement in NF relatively the same as their involvement at baseline.

**Figure 2.**
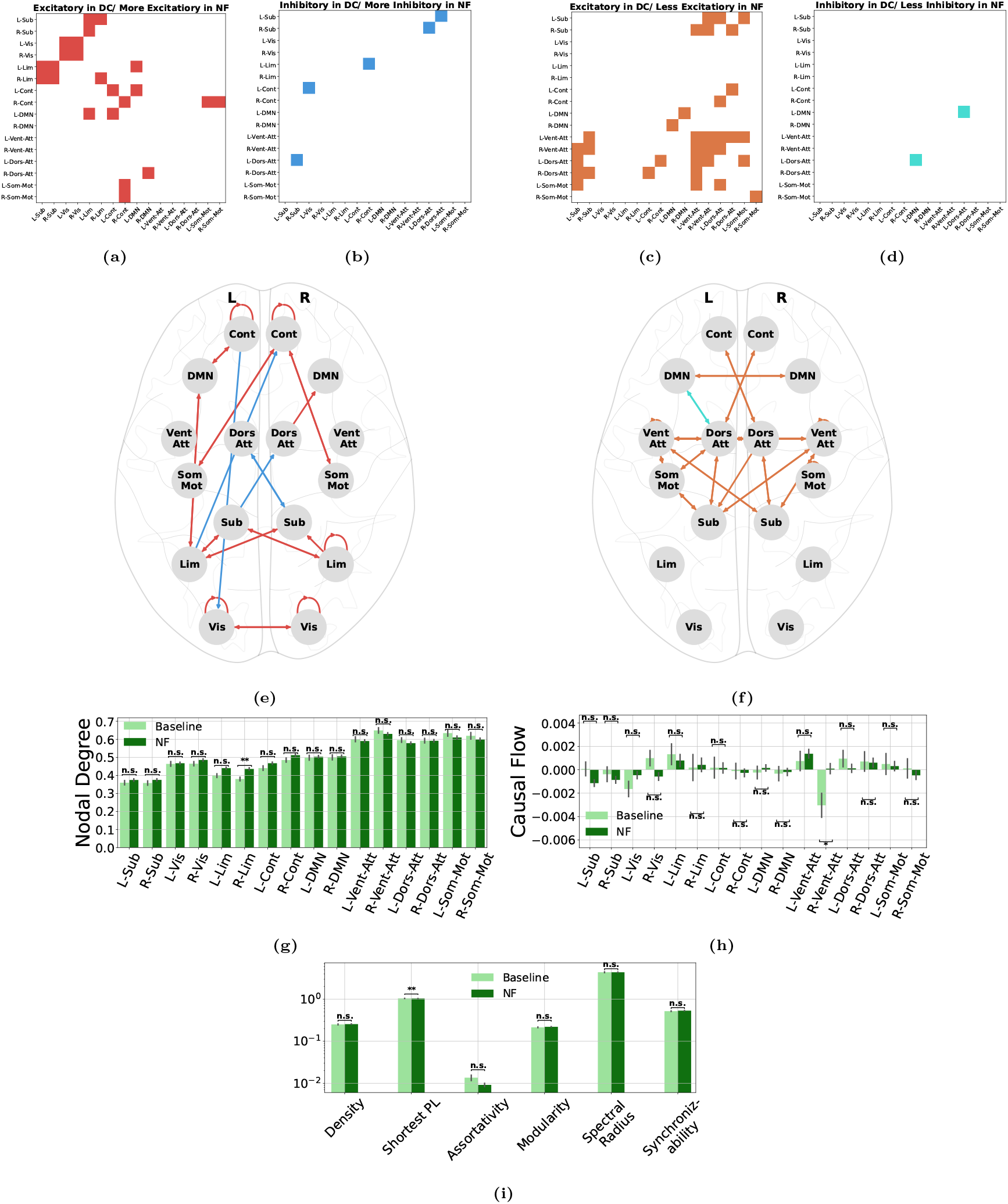
Causal connectivity during NF involves increased engagement of control, limbic, and visual networks and decreased involvement of attention networks. **(a)** Binary heat map showing subnetwork-level causal edges that are excitatory during baseline sessions and became more excitatory during NF. An edge is considered excitatory if its average weight is greater than 0 across all studies and subjects. An edge is considered more excitatory during NF if its weight across all studies and subjects is greater among NF graphs than baseline graphs (*p <* 0.05, two-sided Wilcoxon rank-sum test). Both set of tests are corrected for multiple comparisons using False Discovery Rate (FDR). **(b)** Similar to (a) but for edges that are inhibitory during baseline and become more inhibitory during NF. **(c, d)** Similar to (a, b) but for edges that become less excitatory/inhibitory during NF. **(e)** Color-coded topographic visualization of edges in (a, b). **(f)** Similar to (e) but for edges in (c, d). **(g)** Distribution of nodal degrees across different subnetworks for NF and baseline sessions. **(h)** Similar to (g) but for nodal causal flows. **(i)** Distribution of global network measures during NF and baseline. ^∗^*p <* 0.05, ^∗∗^*p <* 0.01, Wilcoxon rank-sum test, FDR-corrected for multiple comparisons.

Whereas several edges showed a significantly different strength during NF (Figure 2a-f), differences in nodal centralities and global network measures hardly passed the threshold for significance. We observed a significantly higher nodal degree in the right limbic network and a significantly higher causal flow (less sinkness) in the right ventral attention network during NF, while all other nodal degrees and causal flows remained statistically indistinguishable from baseline (Figure 2g,h). Likewise, NF graphs exhibited slightly lower average shortest path length compared to baseline, while their density, assortativity, modularity, spectral radius, and synchornizability remain unchanged (Figure 2i). Thus, in summary, edge-level differences show stronger causal connectivity of control, limbic, and visual networks and weaker connectivity of attention networks during NF compared to baseline, but these differences tend to “average out” when examining more summarized statistics such as nodal centralities or global network measures.

### Causal connectivity within a frontoparietal ‘control hub’ is significantly modulated during NF *and* predicts neurofeedback success

Next, we asked whether subject-specific causal graphs can be used to predict each subject’s success in self-regulation, measured by their trial-by-trial feedback scores. To standardize and ensure comparability of scores across studies, we applied a preprocessing pipeline to trial-by-trial feedback scores and obtained one average standardized score associated with each NF causal graph, as shown in Figure 3a).

**Figure 3.**
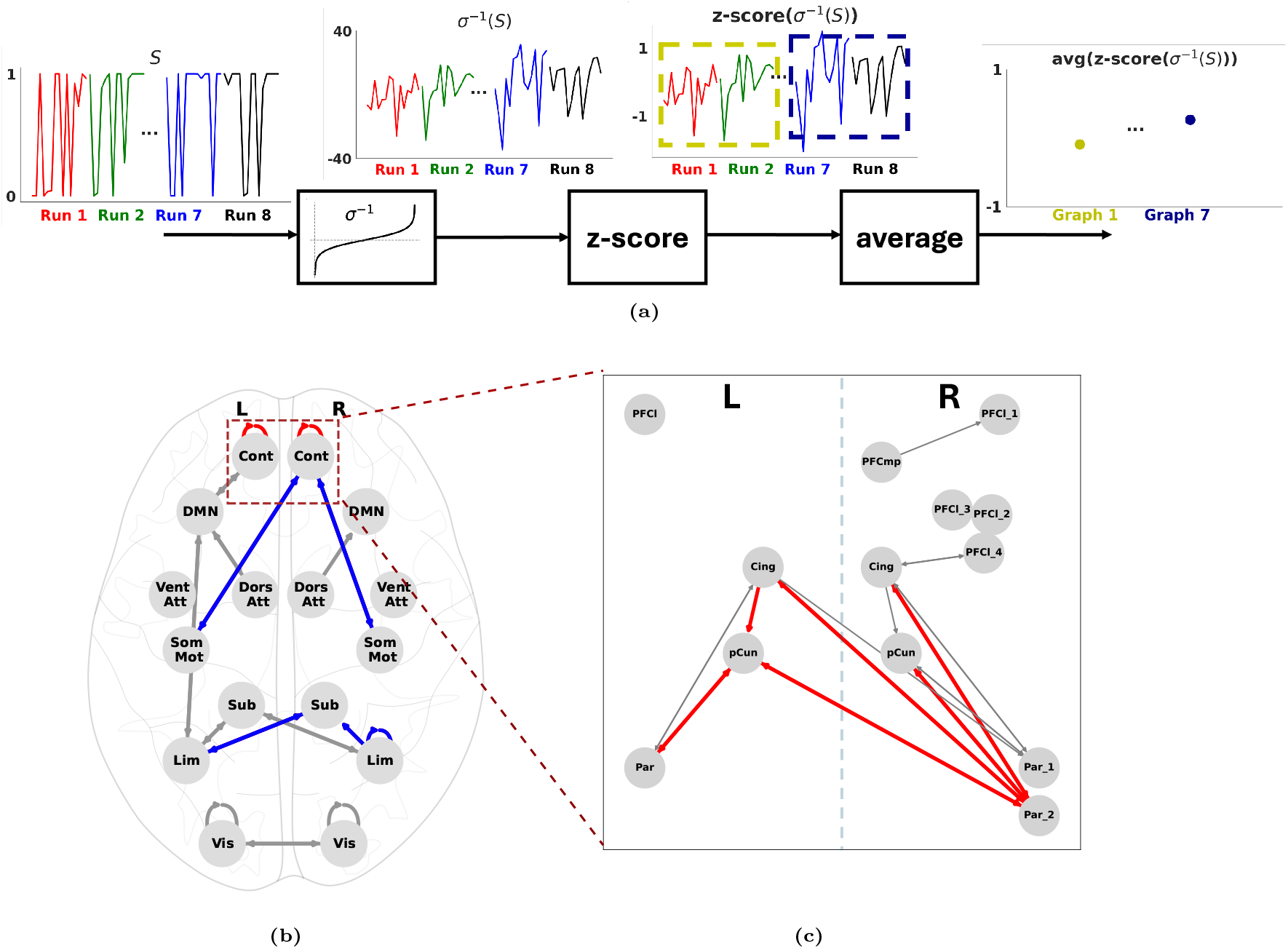
Subjective modulation of a ‘control hub’ predicts successful decoded neurofeedback. **(a)** Preprocessing pipeline for standardizing trial-by-trial feedback scores across DecNef studies. We first passed feedback scores through an inverse sigmoid function to remove the effect of the last sigmoid layer of the logistic regression models used for score computation. This is important since scores from some studies are nearly binary (saturated sigmoids) while scores in other studies are distributed unimodally around 0.5 (Supplementary Figure 3) We then z-scored the results within each study, and averaged them over each pair of consecutive runs to obtain one measure of NF success associated with each causal graph. **(b)** Subnetwork-level causal edges that are significantly stronger in NF sessions compared to baseline and correlate with NF score positively (red), negatively (blue), or insignificantly (gray). Among all edges that are modulated during NF (all depicted edges), only two correlate positively with NF score–within-subnetwork connectivity in the left and right control networks. **(c)** Zoomed-in parcel-level view of the control network. Edge colors have the same meaning as in (b). All red edges connect between bilateral posterior cingulate, precuneus, and lateral posterior parietal cortices, highlighting them as a control hub for decoded neurofeedback.

We then computed the set of edges that were subjectively modulated during NF (NF *>* baseline with FDR-corrected *p <* 0.05, Wilcoxon signed-rank text), and partitioned them based on whether they correlated positively, negatively, or insignificantly with NF success (Figure 3b). Notably, among the 256 subnetwork-level edges, only the two connections within the bilateral control network were modulated during NF *and* correlated positively with NF success. In contrast, connections between right control-bilateral somatomotor and right subcortex-bilateral limbic networks are also subjectively modulated but *negatively* contributed to NF success. The latter may be a reflection of the lack of any motor-related targets in the five analyzed studies, and highlight motor- and emotion-oriented strategies that subjects have been able to consistently recruit, particularly given the lack of any explicit instructions in DecNef, but have been counter-productive for the regulation of specific patterns targeted in each study.

To further investigate the unique contribution of the bilateral control networks to NF success, we zoomed into these networks and repeated the same analysis at the parcel level. As seen from Figure 3c, two main observations are notable. First, we did not observe any edges within the control network that were modulated during NF but correlated negatively with success (no blue edges). Second, and more importantly, all significantly modulated and positively-correlating (red) edges connect between *bilateral posterior cingulate, precuneus, and lateral posterior parietal cortices*. These regions thus function as a hub in decoded neuro-feedback, marking the regions that are not only critical for NF success, but are also actively modulated by subjects during self-regulation.

As seen in Figure 3c, there is noticeable hemispheric asymmetry in the existing nodes and connections within the control network. To ensure that this asymmetry is not due to how the parcels within high-level association cortices are assigned across different brain networks in the Schaefer Atlas, we further extended the parcel-level analysis to include parcels in the default mode network (containing many anatomically-symmetric regions with those of the control network). Interestingly, no additional edges appeared (Supplementary Figure 5), reinforcing the unique role of the aforementioned control hub in successful decoded neurofeedback.

Finally, similar to earlier comparisons, we observed no significant correlations between average neuro-feedback scores and any global measures of causal graphs, including graph density, shortest path length, assortativity, modularity, spectral radius, and synchronizability (Supplementary Figure 4a). Similarly, analyses of nodal centralities, including nodal degree and causal flows for each node, showed no significant associations with neurofeedback scores (see Supplementary Figures 4b and 4c).

**Figure 4.**
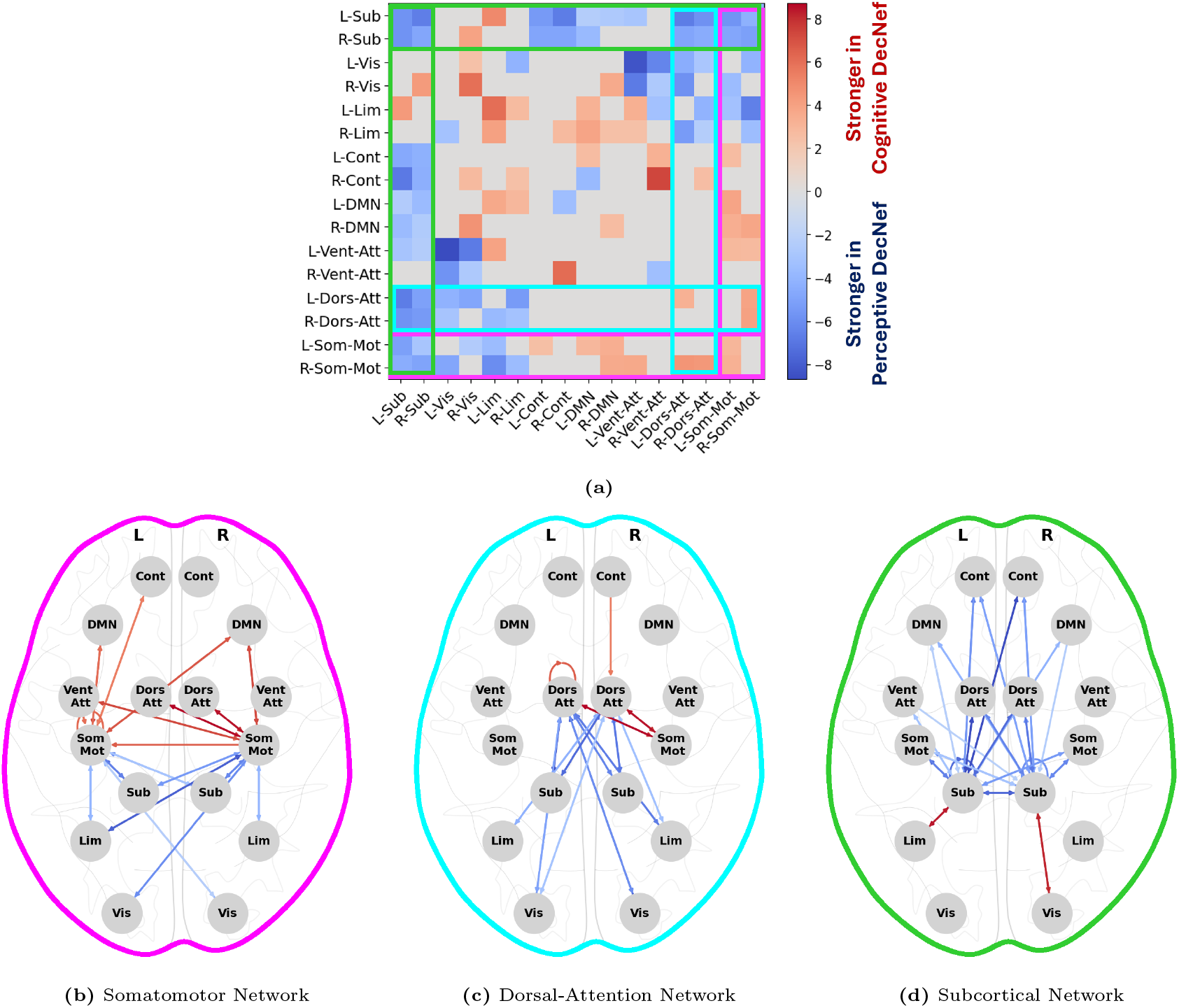
Somatomotor causal connectivity distinctly separates perceptive from cognitive neurofeedback. **(a)** Heatmap illustrating statistically significant differences in causal edge strength between cognitive neurofeedback (studies 1, 4, 5; higher brain functions) and perceptive neurofeedback (studies 2, 3; lower brain functions). Warm (cool) colors indicate stronger connectivity during cognitive (perceptive) neurofeedback (*p <* 0.05, Wilcoxon rank-sum test, FDR corrected). **(b)** Schematic diagram of the subset of edges from (a) associated with the bilateral somatomotor network (color-coded in magenta in (a)). Edges linking the somatomotor network to higher-level networks—control, default mode, and attention—exhibit greater strength during cognitive neurofeedback, whereas edges connecting it to lower-level networks—subcortical, visual, and limbic— show stronger connectivity during perceptive neurofeedback. **(c)** Similar to (b), but for the dorsal attention network. A similar but less prominent pattern is observed compared to that in (b). **(d)** Similar to (b), but for the subcortical network. Here we observe an opposite pattern to those in (b) and (c).

Thus, in summary, subjective modulation of a specific subset of bilateral control network, namely, the posterior cingulate, precuneus, and lateral posterior parietal cortices, is a unique positive predictor of successful self-regulation during decoded neurofeedback, whereas subjective modulation of right control-bilateral somatomotor and right subcortex-bilateral limbic connections negatively correlate with NF success, likely marking common but counter-productive motor- and emotion-oriented regulation strategies.

### Somatomotor causal connectivity distinctly separates perceptive and cognitive neurofeedback

Next, we analyzed how whole-brain causal connectivity differed between DecNef studies that targeted (i.e., sought to modulate cortical patterns related to) cognitive functions and those that targeted perceptive ones. In the five studies in our dataset, two targeted early visual processing (studies 2 and 3) whereas the other three targeted various cognitive functions (face preference in study 1, subjective confidence in study 4, and emotional representation of animals in study 5). We thus grouped studies 2 and 3 into ‘perceptive NF’ and studies 1, 4, and 5 as ‘cognitive NF’. We then tested for the presence of any significant differences in the causal graphs of the two groups of studies. As with previous analyses, we did not find major differences at the level of global network measures (Supplementary Figure 6a) or nodal centralities (Supplementary Figures 6b and 6c). At the edge level, however, several significant differences emerged (Figure 4a). In particular, connections to and from the somatomotor network exhibit a remarkable separation between the two groups of studies: somatomotor connections that are stronger during cognitive NF distinctly link to the control, default mode, and attention networks (hierarchically higher-order), while somatomotor connections that are stronger during perceptive NF distinctly connect to subcortical, visual, and limbic networks (hierarchically lower-order) (Figure 4b). Notably, this strong division appears even though motor regions were not directly targeted in any of the studies. A similar but subtler pattern is observed in the connections to and from the dorsal attention network (Figure 4c), while the opposite pattern exists in connections to and from the subcortical network (Figure 4d). Thus, in summary, these results show the presence of major re-organizations in causal brain connectivity during cognitive and perceptive DecNef, and highlight the somatomotor network as a ‘seed’ that clearly distinguishes between cognitive and perceptive DecNef in its causal connectivity with the rest of the brain.

## Discussion

### Summary

We used whole-brain causal discovery to better understand decoded neurofeedback (DecNef) and its underlying brain mechanisms. Through a meta-study of 5 publicly-available DecNef datasets, we found a particularly salient role for the bilateral control networks, whereby subjects were able to volitionally modulate its internal connections, and this modulation predicted trial-to-trial variability in neurofeedback (NF) success across all studies. This role is unique to the bilateral control network, and specific to a posterior ‘hub’ of the control network consisting of the posterior cingulate, precuneus, and lateral posterior parietal cortices. We further found connections involving the bilateral somatomotor and limbic networks that were volitionally modulated but their modulation predicted NF failure, likely reflecting counter-productive strategies. Given the breadth of our dataset, we were able to further investigate subtle differences between modulation of causal connections in perceptive and cognitive NF. Surprisingly, we found vast differences throughout the brain, including remarkably consistent hierarchical separation in connections to and from the bilateral somatomotor network. Together, our result paint a significantly clearer picture of how human participants engage in decoded neurofeedback, the domain of their volitional modulation, and the effects of such modulation on various DecNef outcomes.

### Network changes during neurofeedback

When combined across all studies and subjects, causal networks during NF sessions showed significantly more variability compared to baseline. This finding may appear to be in contrast to the presence of a shared network mechanisms underlying ‘neurofeedback’, but it is likely due to the open-ended nature of NF training where subjects are not restricted by any explicit instructions or task constraints. Consequently, subjects are free to explore various neural strategies to reach the desired brain state, whereas baseline sessions are structured and task-oriented, requiring participants to complete specific tasks designed to generate a target response. These differences underscore the adaptive nature of NF as an individualized training protocol, particularly for modulating neural circuits underlying complex behaviors.

When comparing NF graphs with baseline, we observed a stronger involvement of the control, limbic, and visual networks, paired with a reduced involvement of attention networks. The strengthening of causal connections involving the control network indicates an increased need for self-regulation and executive control which is crucial for manipulating neural activity. Similarly, the increased involvement of the limbic network likely reflects the emotional and motivational processes that are integral to NF, and the visual network’s increased connectivity with other networks aligns with the exclusive role of vision in delivering feedback in all studies. In contrast, the reduced overall involvement of attention networks, while surprising, may reflect a shift from conventional forms of attention (towards salient exogenous objects or well-defined internal processes) toward freeform internal exploration and unstructured trial and error. Together, these patterns demonstrate subtle but widespread network reconfigurations that characterize the intensive yet creative induction process in decoded neurofeedback.

Though infrequent and subtle, we also observed some global measure and nodal centrality changes in NF compared to baseline. NF graphs displayed a slightly but significantly smaller average path length, suggesting potentially more efficient causal communication pathways during NF. Distinct differences also emerged in nodal centralities of the right limbic and ventral attention subnetworks. NF graphs demonstrated higher nodal degrees in the right limbic network, implying increased connectivity in areas associated with emotional engagement and motivational drive—key factors for maintaining focus and effort during neurofeedback. NF graphs further showed a decrease in causal flow within the right ventral attention network, potentially indicating a shift from external attention processing towards internally directed trial and error.

### The DecNef control hub

A central finding of this meta-study is the key role of posterior cingulate, precuneus, and lateral posterior parietal cortices as a control hub in decoded neurofeedback. The overall control network is widely recognized as a core system supporting high-level cognitive functions such as attention, task management, and goal-directed behavior (Cole et al., 2013; Seeley et al., 2007). It facilitates the integration of information across distributed brain regions, allowing for adaptive responses to dynamic task demands (Cole et al., 2013). Nevertheless, only few regions within the control network (and no regions in the closely-related default mode network) were found to be important for NF success. Existing literature supports posterior cingulate cortex and precuneus as essential regions for orienting attention, maintaining focus, and coordinating between self-referential and externally directed processes (Cavanna and Trimble, 2006; Leech and Sharp, 2014). The posterior parietal cortex complements this by acting as a central node for top-down cognitive control and the rapid reconfiguration of cognitive task sets (Cole et al., 2013; Corbetta et al., 2009; Fang et al., 2024; Lindner et al., 2010; Panidi et al., 2024; Wisniewski et al., 2015). Furthermore, it should be highlighted that increased connectivity *within* this control hub, not between this hub and other regions, was a strong predictor of NF success. In other words, same causal network (as during baseline) between the control hub and the rest of the brain can be sufficient for NF success if connectivity within the control hub itself is properly reorganized. In fact, increased connectivity elsewhere in the network, including those between the control and somatomotor networks, correlated negatively with NF success. From a classical control perspective (Ogata, 2010), this is akin to a feedback control system in which the controller is adaptively modified, while the ‘plant’ and its interconnections to/from the controller remain unchanged.

### Assumptions and limitations

This study has a number of assumptions and limitations, including those implied by the indirect and observational nature of fMRI. As described extensively in (Arab et al., 2025), at the slow temporal resolution of fMRI (TR = 2s here), many causal interactions statistically appear as contemporaneous. This makes it particularly hard to resolve directionalities, particularly in a network with such abundance of feedback as the brain. In addition to the steps taken before in the design of CaLLTiF to address this issue (Arab et al., 2025), we further fine-tuned and adjusted CaLLTiF in this study to compensate for the longer sampling rate of DecNef data (see Methods for details). Another limitation concerns the meta-analysis across studies and subjects. Each DecNef study involved different participants, preventing within-subject comparisons across studies. As such, while our combined dataset offers valuable insights at the group level, it limits our ability to make individual-specific conclusions. Finally, the use of data from decoder construction (DC) sessions as baseline entails limitations. While DC sessions may still be closer to an ideal baseline (identical structure and sensory input as NF without subjects’ effort to self regulate) than the commonly-used resting state baseline, DC sessions include task components that are not present in NF. These task components may partially confound our findings, but their confounding effect is partially mitigated by their variability across studies—that they are partially ‘averaged-out’ in the combination across studies.

### Conclusions

For over two decades, fMRI neurofeedback has remained a promising tool for clinical and cognitive neuroscience; yet neurofeedback studies remain hard to design, understand, and tailor to specific needs and individuals. Using the power of big data, we pinpointed specific subnetworks and whole-brain causal connectivity patterns that address some of the most long-standing questions in neurofeedback research: what neural outcomes humans can and cannot volitionally modulate, what factors determine neurofeedback success, and how volitional modulation varies depending on the target of neurofeedback. Future prospective studies are needed to validate the causality of our findings, particularly in the clinical domain where DecNef has significant potential for empowering individuals to access and manipulate complex circuits underlying psychiatric disorders.

## Material and Methods

### Causal discovery for Large-scale Low-resolution Time-series with Feedback (CaLLTiF)

In this work we used our recently developed causal discovery algorithm CaLLTiF (Arab et al., 2025) to extract causal connectivity graphs from fMRI data. Compared its original form here we slightly modified CaLLTiF to improve its effectiveness with the even slower sampling of the fMRI data in this study (TR = 2s, vs. TR = 0.72s in our earlier work). In CaLLTiF, a causal link is established from a node (parcel) *X*_*i*_ to a node *X*_*j*_ with a lag of *τ* ≥ 0 samples if *X*_*i*_(*t* − *τ* ) is significantly correlated with *X*_*j*_(*t*) after conditioning on all other nodes and their lagged values, ensuring that correlation is not due to a common cause or mediation through other nodes. If *τ* = 0 (contemporaneous effect), a bidirectional feedback connection is placed between *X*_*i*_ and *X*_*j*_, unless at least one of *X*_*i*_ or *X*_*j*_ causes the other also with at least one *τ >* 0, in which case the direction of causality is determined based on the lagged effect(s). However, as discussed in (Arab et al., 2025), lagged effects become exponentially harder to detect with increasing TR and finite samples, even when a statistically significant contemporaneous effect exists, which is itself proof that a lagged effect must have existed. To address this challenge at the slower sampling of 2s, we slightly adjusted CaLLTiF such that for pairs of nodes with a statistically significant contemporaneous effect (detected at the originally-suggested strict significance threshold *α* = 0.0025), we relaxed the threshold of significance on their lagged effects to 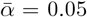. This allows us to uncover weaker lagged connections, and in turn identify additional directional influences that would otherwise be undetectable from purely contemporaneous effects.

As shown in Supplementary Figure 1, increasing the threshold hyper-parameter results in greater asymmetry within the identified causal graphs. For this figure asymmetry measures were computed across all causal graphs (baseline and NF). The maximum asymmetry is observed at a significantly larger 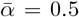. Note that in this case increasing 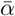 beyond 0.05 is acceptable because this increase is restricted to lagged connections for pairs of nodes between which a contemporaneous effect is already detected (at *α* = 0.0025) and serves as proof that a lagged effect must have existed. Nevertheless, we opted to limit 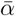 to 0.05 to remain consistent with standard statistical practice.

### Data

We used publicaly available fMRI data from five distinct DecNef studies, published collectively as the DecNef Collection (Cortese et al., 2021). For each participant, data includes a baseline decoder-construction (DC) session for training the machine learning decoderm and several (3 to 10) closed-loop fMRI neural reinforcement sessions. To ensure uniformity and comparability across studies and subjects, data were truncated to 9 subjects per study, 3 NF sessions per subject, 8 and 6 runs per NF and baseline session, respectively, and 152 fMRI volumes per run. We computed one causal graph for each pair of consecutive runs ((NF or baseline), which provided the minimum number of samples needed for CaLLTiF’s partial correlation tests. In total, we obtained 945 causal graphs across NF sessions (5 studies, 9 subjects per study, 3 sessions per subject, 7 graphs per session) and 225 graphs across baseline sessions (5 studies, 9 subjects per study, 1 session per subject, 5 graphs per session).

### Overview of included DecNef studies and targeted neural domains

The five studies included in the DecNef Collection explored DecNef in diverse brain regions and tasks (Cortese et al., 2021). In short, *Study 1* explored facial preference representation in the cingulate cortex (CC), showing that activation patterns within this region could be manipulated to alter preferences for initially neutral faces (Aharon et al., 2001; Chatterjee et al., 2009; Iaria et al., 2008; Said et al., 2011; Shibata et al., 2016). *Study 2* investigated associative learning between orientation and color in early visual areas, demonstrating that DecNef could induce long-term changes in color perception by linking specific visual features such as orientation and color in early visual areas (Amano et al., 2016). *Study 3* examined fear reduction through counter-conditioning in the visual cortex, leveraging DecNef to attenuate conditioned fear responses without explicit awareness (Koizumi et al., 2016). *Study 4* focused on the dissociation between subjective confidence and perceptual accuracy, using DecNef to manipulate confidence without affecting actual performance, challenging the prevailing view that confidence directly reflects perceptual reliability (Cortese et al., 2016; Fleming et al., 2012; Kepecs and Mainen, 2012; Koizumi et al., 2015; Meyniel et al., 2015; Rounis et al., 2010; Simons et al., 2010; Wilimzig et al., 2008). Finally, *Study 5* investigated the unconscious reprogramming of innate fear responses, demonstrating a reduction in physiological fear indicators without conscious exposure to feared stimuli (Guntupalli et al., 2016; Haxby et al., 2011; Taschereau-Dumouchel et al., 2018). Collectively, these studies provide a rich dataset for examining causal brain dynamics across varied neural and behavioral domains, enhancing our understanding of individualized neurofeedback responses.

### Data collection

The fMRI data was acquired using Siemens MAGNETOM Verio and Prisma 3 Tesla MRI scanners. The scanning parameters included a repetition time (TR) of 2000 ms and a voxel size of 3 × 3 × 3.5 mm^3^ (See more details at (Cortese et al., 2021)). All participants across the five studies included in the analysis provided written informed consent. The recruitment procedures and experimental protocols were approved by the institutional review board at the Advanced Telecommunications Research Institute International (ATR, Kyoto, Japan), under the following approval numbers: 14–121, 12–120, 15–181, 14–140, and 16–181. The studies were conducted in accordance with the principles outlined in the Declaration of Helsinki.

### data preprocessing

Raw fMRI data was first preprocessed using standard steps in fMRIPrep (Esteban et al., 2019). Subsequently, we eliminated 9 confounding factors from the time-series data of each voxel. We used Model 3 (9P) in (Ciric et al., 2017) which combines the 6 motion estimates, 2 physiological time series (mean White Matter and mean CSF signals), and the global signal as confounding regressors. Data was then parcellated into 100 cortical regions (Schaefer 100x7 atlas (Schaefer et al., 2018)) and 16 subcortical ones (Melbourne Scale I atlas (Tian et al., 2020)) before being used for causal discovery.

### Aggregation of parcel-level graphs into subnetwork-level graphs

To increase interpretability and reduce the dimensionality of the data, we further aggregated the 116 *×* 116 parcel-level causal graphs into 16 *×* 16 subnetwork-level graphs. The 16 subnetworks consisted of the 7 standard resting-state networks defined by the Yeo-Schaefer atlas (Schaefer et al., 2018; Yeo et al., 2011) (Visual, Somatomotor, Dorsal Attention, Ventral Attention, Limbic, Control, and Default Mode), plus a Subcortical network defined by the Melbourne atlas (Tian et al., 2020), separately in each hemisphere. For any pair of subnetworks, *U* and *V*, the weight of the edge from *U* to *V* was calculated as

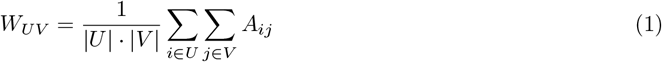

where |*U*| and |*V*| represent the number of parcels in *U* and *V* respectively, and *A*_*ij*_ represents the edge from parcel *i* to parcel *j*. This aggregation was performed in two ways across the analyses in this study: (1) where *A*_*ij*_ is a binary indicator (*A*_*ij*_ = 1 if a significant causal link exists from *i* to *j* and *A*_*ij*_ = 0 otherwise) and *W*_*UV*_ represents the *connection density* from *U* to *V* ; and (2) where *A*_*ij*_ is the magnitude of the partial correlation coefficient from *i* to *j* and *W*_*UV*_ represents the *mean causal strength* flowing from subnetwork *U* to *V* . The former (binary *A*_*ij*_) was used in all calculations of nodal centrality measures and global graph metrics (e.g., Figure 2g-i), whereas the latter (weighted *A*_*ij*_) was used in all edge-wise strength comparisons and score correlations (e.g., Figures 2a-f and 3b).

### Computing nodal degrees and causal flows

The subnetwork-level degrees and causal flows were calculated as

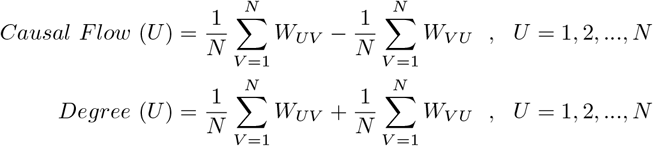

where *W*_*UV*_ is defined in Equation (1) and *N* = 16 is the total number of subnetworks.

### Computing global network measures

Six global network measures were estimated for subnetwork-level graphs, as follows

#### Density

measures the overall probability of the presence of an edge between two nodes. It was computed as the mean *W*_*UV*_ across all edges, where *W*_*UV*_ is defined in Equation (1).

#### Shortest Path Length (PL)

measures how efficiently information can travel across a network (Milgram, 1967; Rubinov and Sporns, 2010). It is computed by calculating the shortest path (i.e., minimum sum of edge weights connecting) between all pairs of nodes using the NetworkX package in Python (Hagberg et al., 2008).

#### Assortativity

measures the tendency of nodes in a network to connect to other nodes that are similar to themselves in some attribute, such as node degree or edge weights. We computed the degree assortativity coefficient as the correlation between the weighted degrees of pairs of connected nodes using the NetworkX package in Python (Hagberg et al., 2008).

#### Modularity

measures the strength of division of a network into communities, quantifying the difference between the observed density of edges within communities and the expected density in a random graph (Newman, 2006). To compute modularity we first converted each subnetwork-level graph into its undirected component (*W* + *W*^⊤^)*/*2 and used the greedy modularity optimization algorithm to detect communities (Newman, 2006).

#### Spectral radius

measures the spread of activity across the network over time, and is given by the largest eigenvalue of the connectivity matrix. A higher spectral radius suggests a stronger, more dominant network structure, with greater potential for synchronization and transitions into excited state (Meghanathan, 2014; van Dam and Kooij, 2007; Wang et al., 2015, 2003).

#### Synchronizability

measures how easily a network can synchronize its components, reflecting the stability and collective behavior of the network when nodes attempt to synchronize (Arenas et al., 2008; Tang et al., 2014). After calculating the Laplacian matrix of each subnetwork-level graph using the NetworkX package in Python (Hagberg et al., 2008), we computed synchronizability

## Computing

All the computations reported in this study were performed on a Lenovo P620 workstation with AMD 3970X 32-Core processor, Nvidia GeForce RTX 2080 GPU, and 512GB of RAM.

## Additional Information

## Author contributions

EN designed and supervised the study; FA performed all analyses; AG assisted with the design and interpretations of the causal discovery algorithm; HJ and MAKP assisted in the analyses of human fMRI data; FA and EN drafted and all authors edited the manuscript.

## Acknowledgments

The research conducted in this study was partially supported by NSF Award #2239654 to EN, by the Canadian Institute for Advanced Research (fellowship awarded to MAKP), and by the Air Force Office of Scientific Research under award number FA9550-20-1-0106 (to MAKP).

## Data availability statement

All the fMRI data used in this work is publicly available. The fMRI data from DecNef studies can be accessed upon request (Cortese et al., 2021).

## Code availability statement

The Python code for this study is available upon request and will be publicly released upon publication.

