## Supplementary Figures for "Whole-brain causal connectivity during decoded neurofeedback: a meta-study"

### Supplementary Material for “Whole-brain causal connectivity during decoded neurofeedback: a meta study”

[1]*Department of Radiology and Biomedical Imaging, University of California, San Francisco, USA*
[2]*Department of Mathematics and Statistics, Boston University, USA*
[3]*Department of Psychiatry and Psychotherapy, Phillips University of Marburg, Germany*
[4]*Department of Cognitive Sciences, University of California, Irvine, USA*
[5]*Center for the Neurobiology of Learning & Memory, University of California, Irvine, USA*
[6]*Program in Brain, Mind, & Consciousness, Canadian Institute for Advanced Research, Canada*
[7]*Department of Mechanical Engineering, University of California, Riverside, USA*
[8]*Department of Bioengineering, University of California, Riverside, USA*
[9]*Neuroscience Graduate Program, University of California, Riverside, USA*
[10]*Department of Electrical and Computer Engineering, University of California, Riverside, USA*

### Supplementary Figures

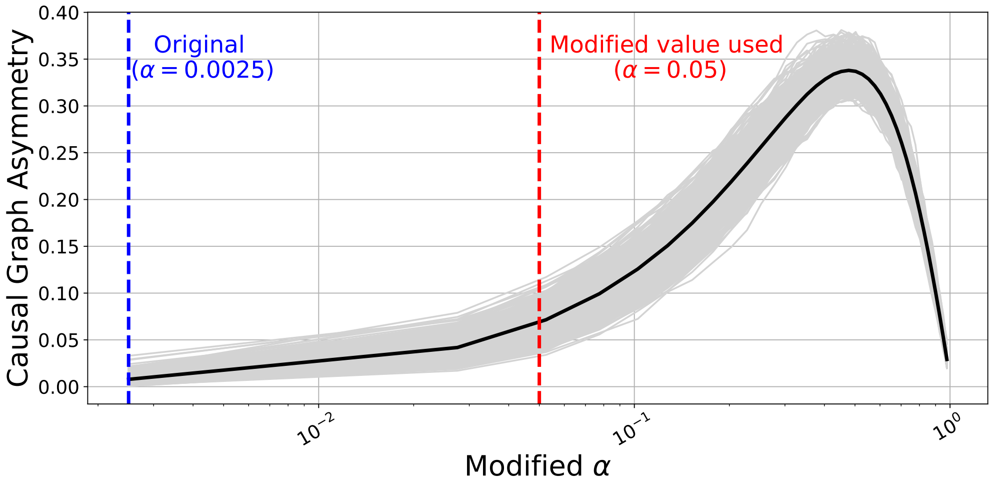


**Supplementary Figure 1: The asymmetry of causal graphs computed by the adjusted CaLLTiF algorithm.** As described in the main text, the type I error bound was adjusted for detecting lagged partial correlations between pairs of nodes for which a contemporaneous effect has already been found. Graph asymmetry initially increases with increased detection of lagged edges (increased modified $\alpha$) as directionality can only be determined for lagged edges. We thus monitored graph asymmetry as a measure of the effect of increasing modified $\alpha$. Asymmetry for a graph represented by matrix $A$ is computed as $\frac{\|A-A^T\|_1}{\|A+A^T\|_1}$. The asymmetry reaches a peak around a threshold value of 0.5. After the peak further increases in the threshold lead to a decrease in asymmetry as the graph becomes fully symmetric at the limit of modified $\alpha = 1$.

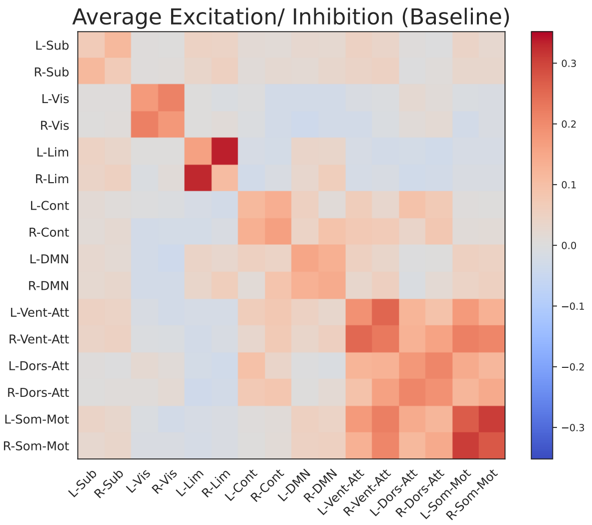


**(a)**

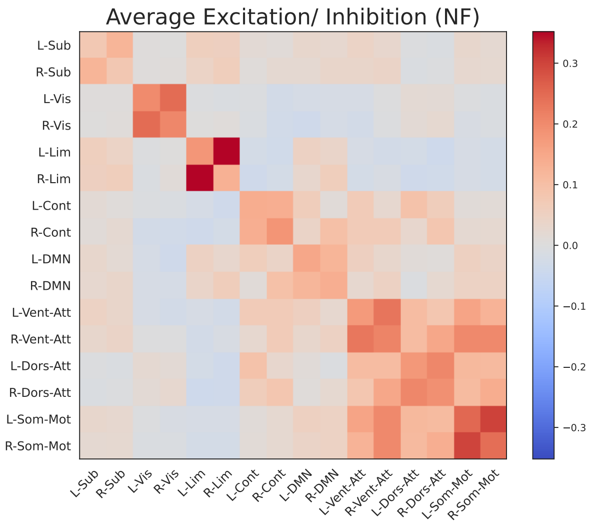


**(b)**

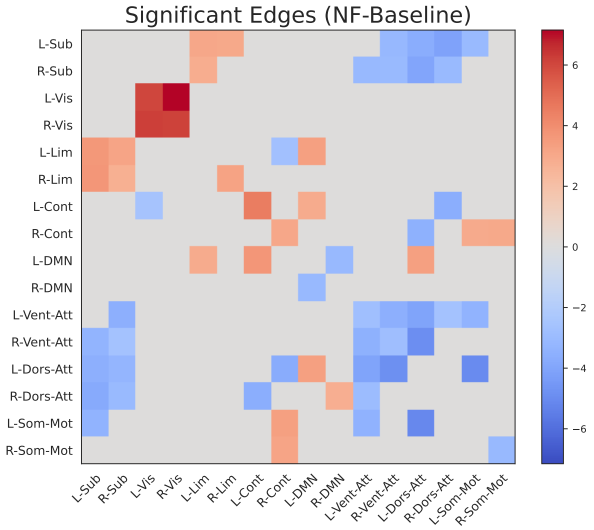


**(c)**

**Supplementary Figure 2: NF and baseline whole-brain causal graphs mainly consist of excitatory connections. (a)** Average causal graph (among all studies, subjects, sessions, and runs) computed from partial correlation maps of baseline sessions. **(b)** Average causal graph (among all studies, subjects, sessions, and runs) computed from partial correlation maps of NF sessions. Since edge weights in each causal graph are derived from partial correlations, their sign can provide insights into whether the respective effective connections have a net excitatory or net inhibitory effect. Most causal effects in baseline and NF sessions are excitatory. **(c)** Significant differences in NF sessions compared to baseline (difference of edge weights multiplied by $p < 0.05$, Wilcoxon rank-sum test, FDR-corrected for multiple comparisons).

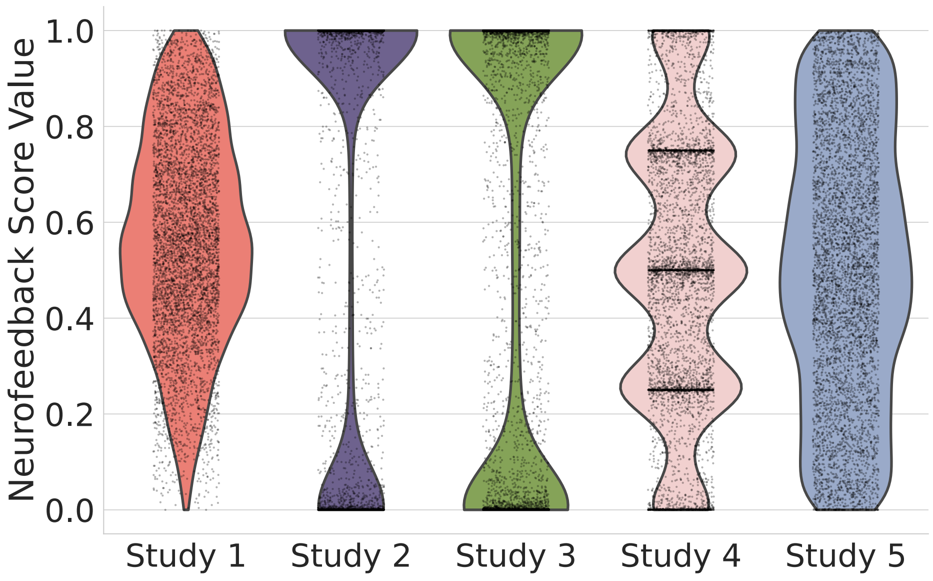


**Supplementary Figure 3: Distribution of neurofeedback induction scores across five DecNef studies.** Violin plots show the distribution of decoder output scores (0–1) aggregated across all subjects and trials for each study. Studies exhibit largely distinct distributions, particularly 2 and 3 vs. 1, 4, and 5. As noted in the main text, this demonstrates the saturation of logistic regression final sigmoids in studies 2 and 3 and prevents/biases comparisons of raw scores across studies.

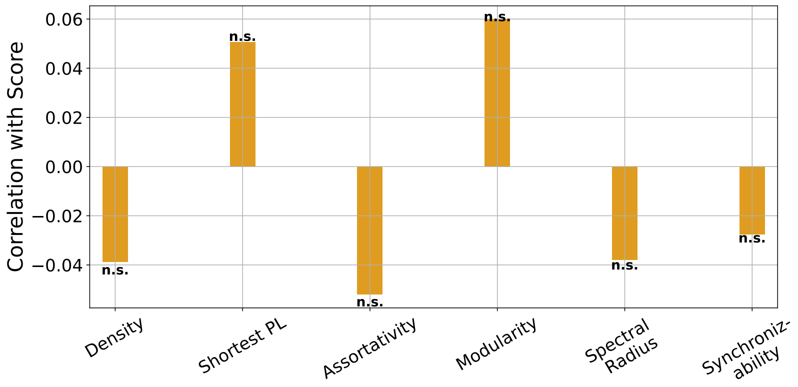


**(a)**

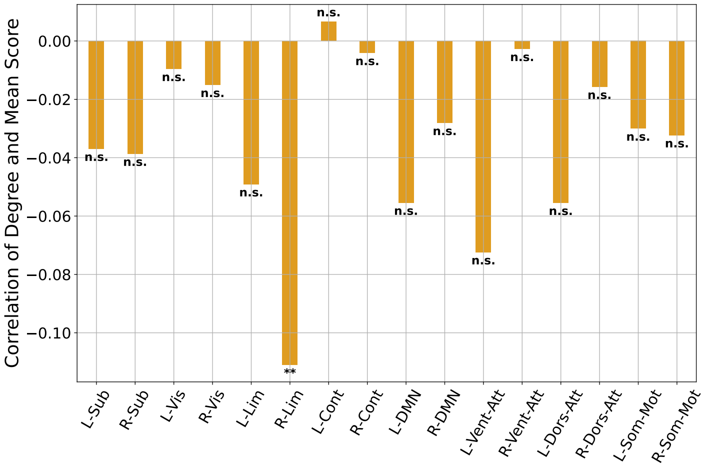


**(b)**

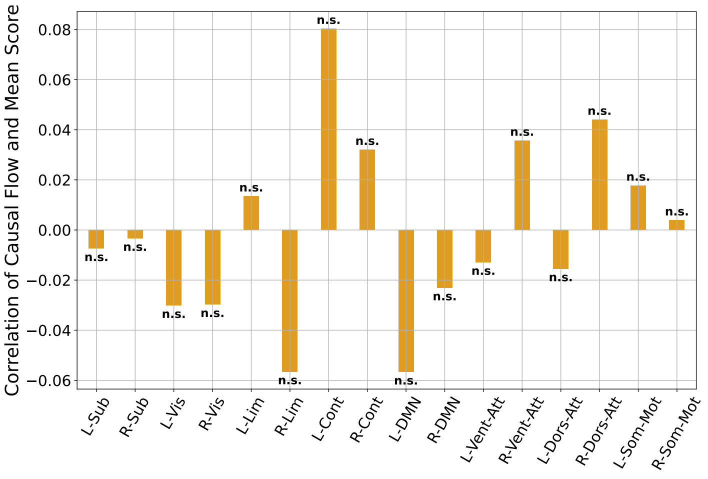


**(c)**

**Supplementary Figure 4: Lack of significant correlations between average NF scores and global measures or nodal centralities of NF causal graphs.** **(a)** Correlations between average NF scores and global network measures. **(b)** Correlations between average NF scores and nodal degrees. **(c)** Correlations between average NF scores and nodal causal flows. All statistical comparisons are performed using Student's t-test at $\alpha = 0.05$ and FDR corrected for multiple comparisons.

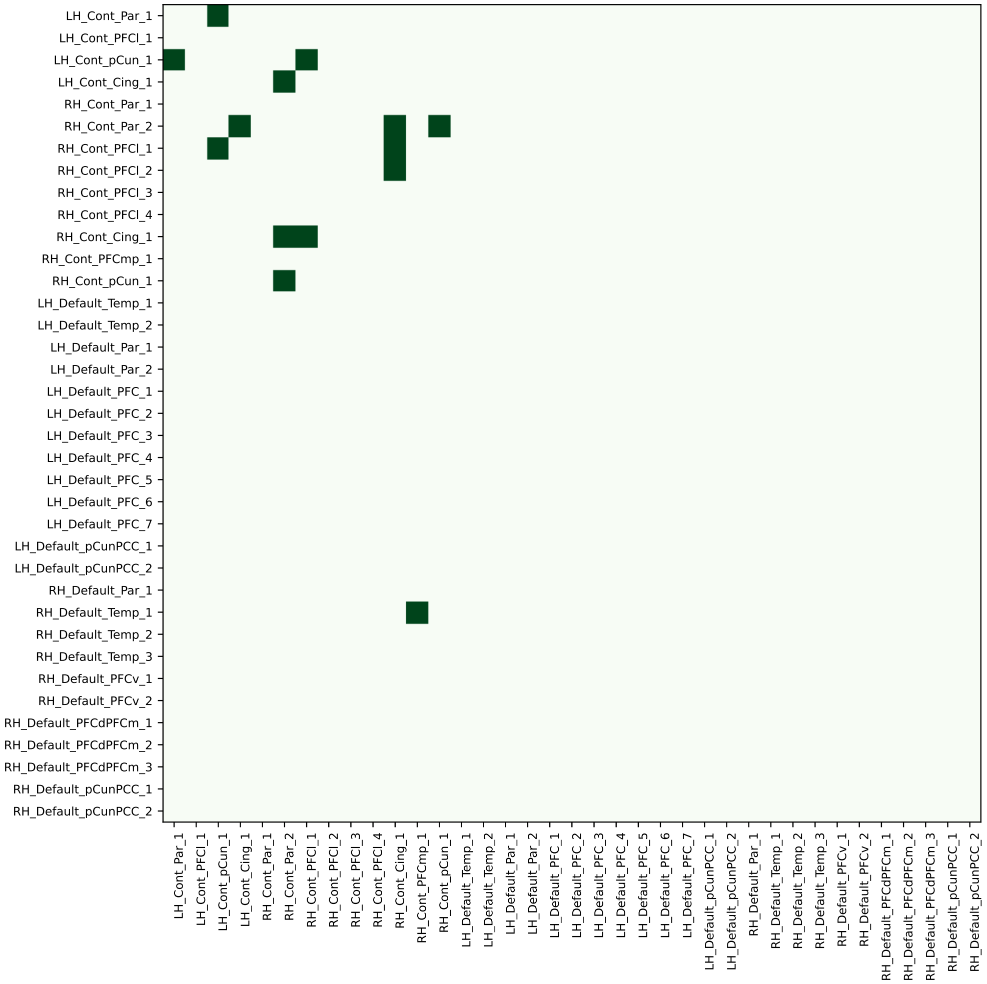


**Supplementary Figure 5: Hemispheric asymmetry in correlations with NF scores observed within the control network is likely an intrinsic characteristic rather than a result of network parcellation.** Dark cells indicate edges that were stronger in NF compared to baseline and correlate positively with NF scores, including all parcels (and their connections) from the control and default mode networks (DMN). As noted in the main text, we specifically examined the DMN because in the Schaefer parcellation it includes several parcels at anatomically mirrored locations to those in the control network.

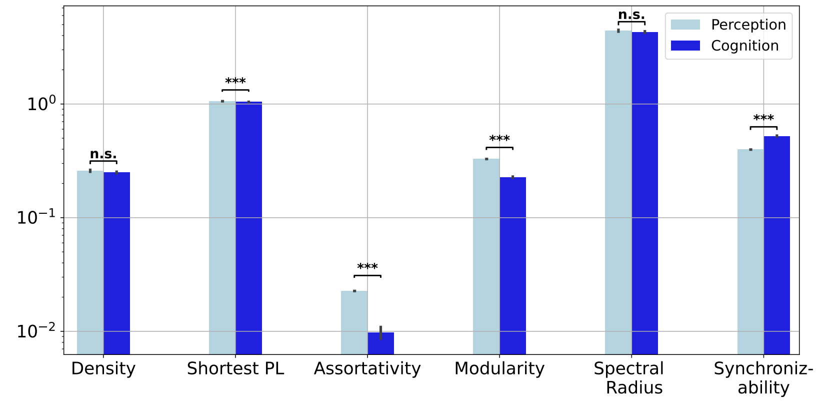


**(a)**

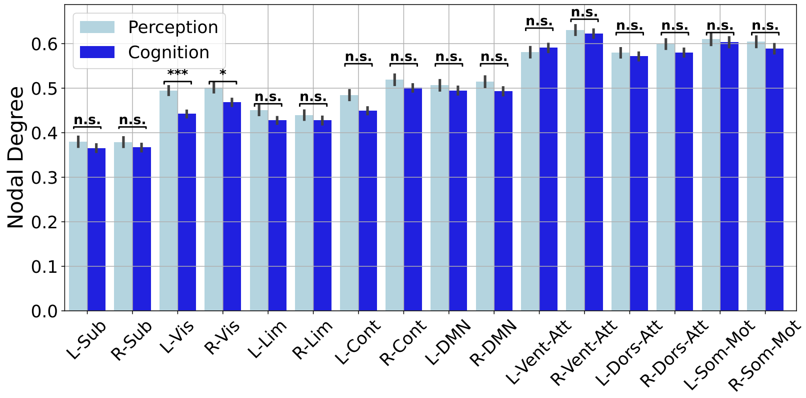


**(b)**

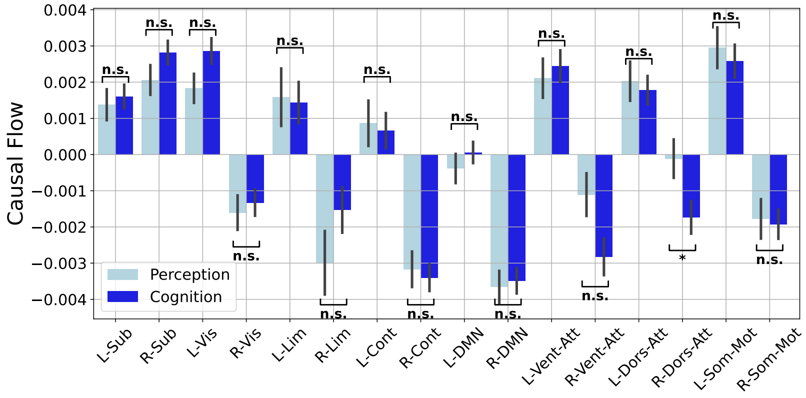


**(c)**

**Supplementary Figure 6: Lack of major differences at the level of global network measures or nodal centralities between cognitive and perceptive NF. (a)** Distribution of global network measures in cognitive and perceptive NF causal graphs. **(b,c)** Similar to (a) but for nodal degrees and causal flows. All statistical comparisons are parformed using Wilcoxon rank-sum test at $\alpha = 0.05$ and FDR corrected for multiple comparisons.
